# Backtracking Metabolic Dynamics in Single Cells Predicts Bacterial Replication in Human Macrophages

**DOI:** 10.1101/2024.10.28.620606

**Authors:** Mariatou Dramé, Dmitry Ershov, Jessica E. Martyn, Jean-Yves Tinevez, Carmen Buchrieser, Pedro Escoll

**Affiliations:** Institut Pasteur, Université Paris Cité, Biologie des Bactéries Intracellulaires, Département de Microbiologie, F-75015 Paris, France; Institut Pasteur, Université Paris Cité, Image Analysis Hub, F-75015 Paris, France; Institut Pasteur, Université Paris Cité, Bioinformatics and Biostatistics Hub, F-75015 Paris, France

**Keywords:** *Legionella pneumophila*, intracellular bacteria, macrophages, prediction, mitochondria, mitochondrial membrane potential, reactive oxygen species, machine learning

## Abstract

Accurately tracking dynamic state transitions is crucial for modeling and predicting biological outcomes, as it captures heterogeneity of cellular responses. To build a model to predict bacterial infection in single cells, we have monitored in parallel infection progression and metabolic parameters in thousands of human primary macrophages infected with the intracellular pathogen *Legionella pneumophila*. By combining livecell imaging with a novel tool for classifying cells based on infection outcomes, we were able to trace the specific evolution of metabolic parameters linked to distinct outcomes, such as bacterial replication or cell death. Our findings revealed that early changes in mitochondrial membrane potential (Δψm) and in the production of mitochondrial Reactive Oxygen Species (mROS) are associated with macrophages that will later support bacterial growth. We used these data to train an explainable machine-learning model and achieved 83% accuracy in predicting *L. pneumophila* replication in single, infected cells before bacterial replication starts. Our results highlight backtracking as a valuable tool to gain new insights in host-pathogen interactions and identify early mitochondrial alterations as key predictive markers of success of bacterial infection.

## Introduction

The response of cells, such as macrophages to bacterial infection has for long time been studied by analyzing the bulk output of an ensemble of cells. However, individual cells can exhibit heterogeneous responses to infection. Recent years have witnessed exciting technological and bioinformatics advances that have enabled the detailed profiling of single cells allowing to explore the heterogeneity of the cellular response to infection. Furthermore, the increased accessibility and affordability of high-throughput technologies now makes it possible to collect large datasets that capture the heterogeneity of individual cell responses and explore the transitions between different cellular states. High-throughput imaging and analyses of living, single cells, in particular, facilitate precise measurements of the dynamic state transitions across thousands of cells simultaneously (1). Since cells remain alive throughout the observation period, live-cell imaging uniquely allows for the continuous tracking of the behavior of each individual cell and the monitoring of transitions in individual cells. This approach allows capturing the heterogeneity of individual responses within cell populations. To predict biological outcomes in such complex systems, it is essential to computationally process state transitions in these systems (1, 2), which is crucial to train models aiming to predict biological outcomes.

Predicting the progression of bacterial infection, at the single cell scale, could help to identify early markers of infection, develop personalized therapeutic strategies, and enhance our understanding of the cellular mechanisms that determine infection outcomes (3–6). To train a predictive model, the heterogeneity of individual responses in bacterial and host cell populations should be captured in large datasets that can be then decoded by the model. Here we chose *Legionella pneumophila*, an intracellular growing bacterial pathogen, and infection of primary human macrophages as our model, as infection by intracellular bacteria might represent a valuable model to capture this heterogeneity in hostpathogen interactions. Intracellular bacteria invade and replicate in host cells where they create a unique niche in each infected cell that can be easily tracked and analyzed (7). Compared to the complex spread of extracellular bacteria throughout an organism, studying infection by intracellular bacteria simplifies the tracking of dynamic state transitions of both the bacterial and the host individual, living cells, as these dynamics can be observed within the confines of the infected cell. Macrophages are innate immune cells at the frontline defense against pathogens. They are present in all tissues and show a large diversity in terms of anatomy, transcriptional profiles and functional capacities (8, 9). Although they are specifically armed to fight bacterial infections, many intracellular bacteria can infect and replicate within macrophages during human disease (10–12). Examples are *Mycobacterium tuberculosis* (13), *Salmonella enterica* (14), *Chlamydia trachomatis* (15), or *Legionella pneumophila* (16). These bacteria are also known to manipulate the metabolism of infected macrophages to their own advantage (17–23).

A critical challenge for these intracellular bacteria is to obtain the specific nutrients to grow inside infected cells. As these need to be obtained from the infected cell, modulating the metabolism of host cells to fuel pathogen growth during infection has been shown to be a virulence strategy employed by intracellular bacteria to grow intracellularly (24– 26). Concomitantly, host cells have also evolved mechanisms to detect pathogen-induced alterations in their metabolism and trigger cell-autonomous immune responses against infection (25, 27–31). Thus, the specific evolution at the single cell scale of these metabolic interactions between intracellular bacteria and the host cells may have an important role dictating the outcome of an infection event.

In the present study, we aimed to build a model able to predict the outcome of *L. pneumophila* infection in human macrophages. We used primary macrophages because cancer cell lines exhibit altered metabolism, which can influence the replication of intracellular bacteria (32–34). Indeed, during infection of humans, *L. pneumophila* replicates within lung alveolar macrophages, causing a severe form of pneumonia known as Legionnaires’ disease (16, 35). Infection is facilitated by a Type IV secretion system (T4SS), which translocates over 330 bacterial effectors from the *Legionella*containing vacuole (LCV) into the host cell cytosol (35–37). These effectors play critical roles in manipulating various host cell functions to promote bacterial survival and replication. Interestingly, among those several T4SS effectors target mitochondrial functions, which modify the metabolic state of infected macrophages to help support bacterial growth (38– 42)

Thus, here we analyzed the dynamics of mitochondrial and cellular parameters in infected cells at the single cell level. We developed an innovative imaging pipeline using spinning-disk high-throughput confocal microscopy to conduct time-lapse experiments on thousands of *L. pneumophila*infected human macrophages, allowing us to track individual cells and monitor the progression of bacterial infection while continuously monitoring metabolic parameters. To correlate metabolic responses with infection outcomes at the singlecell scale, we developed BATLI (Backtracking Analysis of Time-Lapse Images), a software tool for retrospective analysis of individual infected cells. BATLI allowed us to categorize cells based on outcomes like bacterial replication or cell death and identify specific metabolic alterations occurring already before the onset of bacterial replication that are linked to these outcomes.

We found that changes in mitochondrial functions, occurring prior to the onset of *L. pneumophila* replication, are predominantly measured in macrophages that will later support intracellular, bacterial growth. We trained an explainable machine-learning model, which predicted already at 5 hours after infection with 83% accuracy, whether an infected cell would later support or restrict bacterial replication. Our results suggest that mitochondrial functions are predictive biomarkers of infection progress and highlight the potential of analyzing metabolic dynamics at the single cell scale to accurately predict infection outcomes.

## Results

### *Single-cell analysis of infected macrophages reveals that replication of* L. pneumophila occurs only in a small subset of permissive host cells

To investigate whether each infected macrophage permits *L. pneumophila* replication or not, we infected human monocyte-derived macrophages (hMDMs) at a multiplicity of infection (MOI) of 10 with either wild-type *L. pneumophila* strain Paris (Lpp-WT) or an avirulent mutant lacking a functional T4SS necessary for effector translocation (Lpp-Δ*dotA*). This mutant strain, unable to replicate inside macrophages, served as a control (43, 44). Both bacterial strains carried the pNT28-GFP plasmid (45), constitutively expressing GFP (hereafter designated as LppWT-GFP and Lpp-Δ*dotA*-GFP, respectively). We performed infection assays in 384-well microplates, to simultaneously examine multiple experimental conditions in hMDMs obtained from the same donor (46). For each condition at least four technical replicates were conducted. To track individual cells, we stained hMDMs with Hoechst and Cell Tracker Blue dyes (**Table 1**), facilitating the detection of nuclei and cytoplasm using a single microscope channel. Hoechst provided a strong nuclear signal, while Cell Tracker Blue gave a dim signal in the cytoplasm (**Figure 1A)**. To capture the dynamics of infection, we acquired time-lapse images of thousands of live hMDMs infected with Lpp-WT-GFP or LppΔ*dotA*-GFP at hourly intervals up to 18 hours post-infection (hpi). This approach enabled us to observe the entire infection cycle of *L. pneumophila* in human cells within a single session.

**Table 1.** List of dyes used to measure cellular parameters in living, single cells. * Channel (excitation/emmision, in nm): Blue = 375/435-480, Green = 488/500-550, Red = 561/570-630, Far Red = 640/650-760. N/A = Non-Applicable.

| Measured Parameter | Image Analysis Script output | Dye | Channel* | Concentration |
| --- | --- | --- | --- | --- |
| Nuclei size | Area of Nucleus ($\mu\text{m}^2$) | Hoechst 33342 | Blue | 200 ng/mL |
| Nuclei condensation | Mean Fluorescence Intensity in the Nucleus | Hoechst 33342 | Blue | 200 ng/mL |
| Nuclei morphology | Nuclei Roundness | Hoechst 33342 | Blue | 200 ng/mL |
| Cell size | Area of Cytoplasm ($\mu\text{m}^2$) | Cell Tracker Blue | Blue | 25 $\mu\text{M}$ |
| Cell morphology | Cytoplasm Roundness | Cell Tracker Blue | Blue | 25 $\mu\text{M}$ |
| Early cell death | Mean Fluorescence Intensity in the Cytoplasm | Annexin V | Red, Far Red | 1/100 |
| Late cell death | Mean Fluorescence Intensity in the Nucleus | Sytox | Far Red | 0.25 $\mu\text{M}$ |
| Mitochondrial membrane potential | SD/Mean of Fluorescence Intensity in the Cytoplasm | TMRM | Red | 25 nM |
| Mitochondrial mass | Sum of the area of intracellular structures (spots) ($\mu\text{m}^2$) | MitoTracker | Red, Far Red | 50 nM |
| Mitochondrial morphology | SER Edge and SER Ridge | MitoTracker | Red, Far Red | 50 nM |
| Cellular Reactive Oxygen Species | SD/Mean of Fluorescence Intensity in the Cytoplasm | CellROX | Red, Far Red | 5 $\mu\text{M}$ |
| Bacterial Vacuole | Sum of the area of intracellular structures (spots) ($\mu\text{m}^2$) | GFP | Green | N/A |

**Figure 1.**
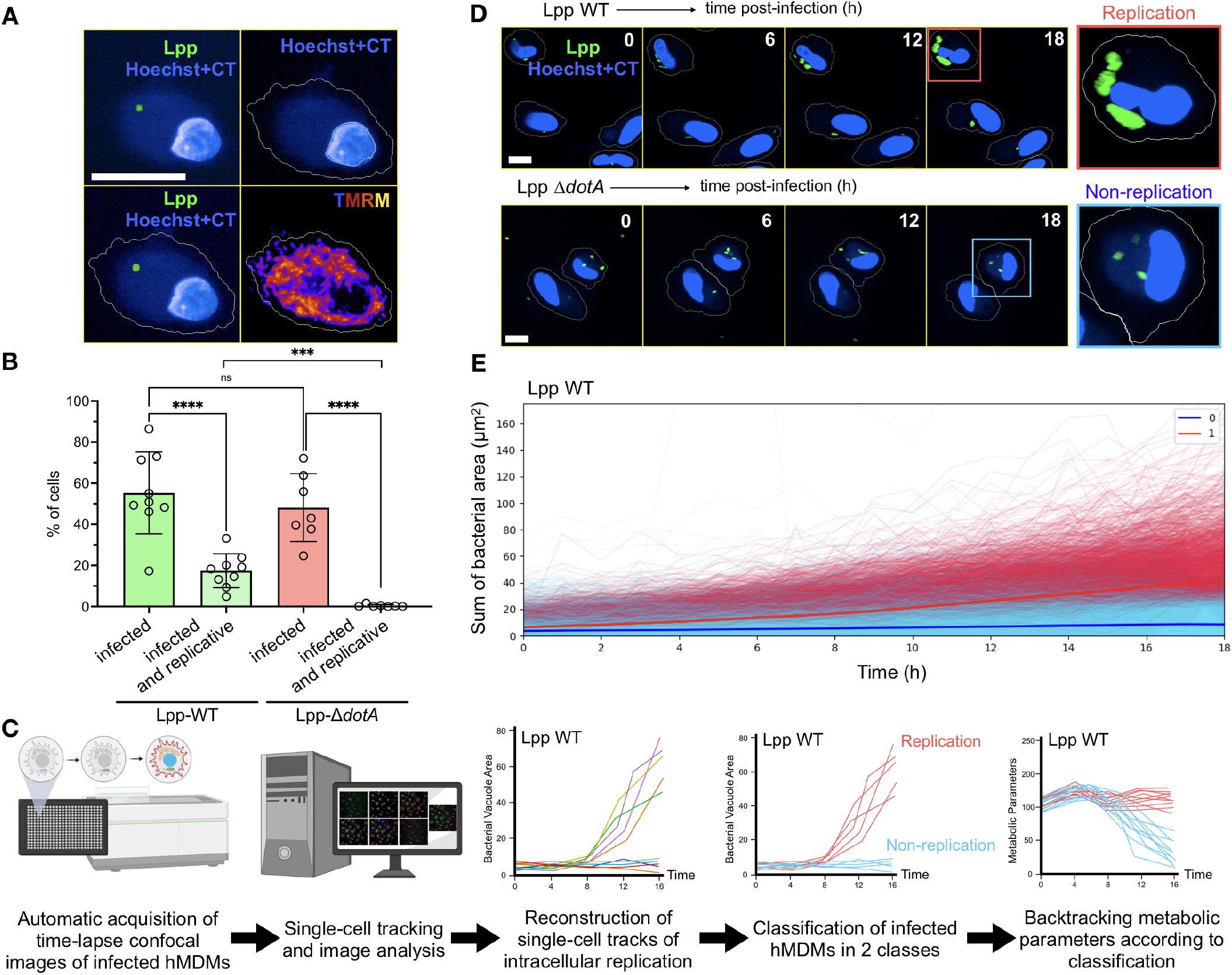
Analyses of single-cell trajectories of macrophages during infection allows their classification as supporting or restricting intracellular replication of *L. pneumophila*. (**A**) Confocal images of a hMDM infected with *L. pneumophila* strain Paris (Lpp) wild-type (WT). GFP-expressing Lpp-WT is shown in green, while Hoechst-stained nuclei and Cell Tracker Blue (CT)-stained cytoplasm are shown in blue. These signals were used to segment nuclear, cytoplasmic and bacterial compartments in each cell by automatic image analysis (white lines). Staining with TMRM dye to measure mitochondrial membrane potential (Δψm) is shown in pseudo-color (hot lava) and was measured in the cytoplasm of each infected cell. Bar = 25 µm. (**B**) hMDMs were infected with Lpp-WT or the T4SS-deficient strain Lpp-Δ*dotA* (non-replicative), and stained with Hoechst+CT to track their nuclei and cytoplasm during 18 hours post-infection (hpi). We defined bacterial replication as occurring in a cell if: (bacterial area at t=18 hpi) (bacterial area at t=0 hpi) > 15 µm^2^. The percentage of infected cells at t = 0 hpi and the percentage of infected cells with replicative vacuoles at t = 18 hpi are shown for both strains (ns = non-significant, ***=p<0.001, ****= p<0.0001, ordinary one-way ANOVA). (**C**) Schematic representation of the analysis of macrophage single-cell tracks during infection and bacterial replication-dependent classification. hMDMs were infected by either the Lpp-WT or the Lpp-Δ*dotA* strain, stained with Hoechst+CT to track their nuclei+cytoplasm and dyes for living cells to measure metabolic parameters (see Table 1). After automatic acquisition of time-lapse confocal images of hMDMs and computational image analysis for replication dynamics, a threshold for replication was set using the formula used in (B). Dynamics of hMDMs infected by the non-replicative Lpp-Δ*dotA* strain was used as control. BATLI software was then used to reconstruct single-cell trajectories and classify cells according to cell fate. Cellular parameters were then backtracked in every single, infected cell and plotted according to their classification. (**D**) Confocal images of infected hMDMs during the infection course, from 0 to 18 hpi. GFP-expressing Lpp are shown in green and Hoechst+CT-stained cells to segment nuclei and cytoplasm are in blue. Red squares highlight cells containing replicative vacuoles while blue squares show cells with non-replicative vacuoles. Bar = 25 µm. (**E**) Single-cell tracks of bacterial vacuole sizes in µm^2^ for Lpp WT-infected hMDMs during infection course (0 to 18 hpi), each single line represents one cell. Single-cell trajectories classified by BATLI as hMDMs with replicative vacuoles are shown in red, while single-cell trajectories classified as hMDMs with non-replicative vacuoles are shown in blue.

Using our image analysis pipeline, we identified and selected individual, infected cells in each time-lapse experiment and monitored bacterial replication within each macrophage by measuring the area occupied by intracellular *L. pneumophila* (GFP fluorescence area, referred to as bacterial area, **Figure 1A)**. To minimize the bias due to variations in bacterial entry we need to define bacterial replication as an increment in occupied area at late time-points (t=18 hpi) compared to early time-points (t=0 hpi). To do this, we first measured the area occupied by a single, intracellular bacterium in 50 cells, which gave values that ranged between 0.6 and 2.6 µm^2^ (**Figure S1A)**. Based on these values we defined bacterial replication as occurring in a cell if: (bacterial area at t=18 hpi) (bacterial area at t=0 hpi) was > 15 µm^2^, which corresponds to a bacterial population duplicating at least three times in a cell.

All together, we examined 76,995 single Lpp-WT-GFPinfected cells across 9 independent experiments and 34,717 single Lpp-Δ*dotA*-GFP-infected cells across 7 independent experiments. At the initial time point (t = 0), 55 ± 19% of hMDMs were infected by Lpp-WT-GFP, while 48 ± 15% were infected by the Lpp-Δ*dotA*-GFP strain. The infection rates were not significantly different, suggesting that invasion of hMDMs primarily depends on the phagocytic capacity of the host cell rather than the presence of T4SS effectors (**Figure 1B)**.

At 18 hpi, 17 ± 8% of the Lpp-WT-GFP-infected macrophages bacterial replication was observed, while only 0.4 ± 0.5% of Lpp-Δ*dotA*-GFP-infected macrophages showed signs of bacterial replication (p<0.005). This low replication rate for the T4SS mutant is consistent with its inability to establish an LCV and validates our criteria for defining bacterial replication. Our results indicate that *L. pneumophila* replication in hMDMs is relatively inefficient, with exponential growth occurring in only a small sub-set of permissive macrophages. This suggests that some heterogeneity at the host-pathogen interface significantly influences the replication of *L. pneumophila* in human primary macrophages.

### *Backtracking mitochondrial functions in macrophages revealed that early alterations occur preferentially in macrophages that support the replication of* L. pneumophila

To capture the heterogeneity of cellular responses in infected macrophages and analyze its impact on the intracellular replication of bacteria, we tracked phenotypes of single cells over time, analyzed host and bacterial dynamics within each infected cell, and determined if cellular parameters correlated with bacterial replication. To facilitate these analyses, we developed an open-source software named BATLI (Backtracking Analysis of Time-Lapse Images). Using BATLI we could reconstruct the history of each cell from time-lapse experiments and perform retrospective analyses of dynamic measurements of fluorescence and morphological properties. Time-lapse images were recorded from a large number of cells using automated confocal microscopy and microtiter plates (384-well plates in this study). Infected cells and cellular compartments, such as the nucleus, mitochondria, cytoplasm, and bacterial vacuole, were automatically detected through segmentation using signals from different channels. Then we used BATLI to classify single-cell trajectories at the end of the infection into two categories: macrophages showing bacterial replication and those not showing bacterial replication (**Figure 1C)**.

We reconstructed and analyzed the trajectories of 20,026 human macrophages infected with the WT strain across nine independent experiments. As control, the avirulent Δ*dotA* mutant strain was used to validate the threshold for bacterial replication used in our analysis (**Figure 1D** and **S1B-D**). Aggregating data from all experiments, we observed that 21% of WT-infected hMDMs exhibited bacterial replication. This proportion aligns with our earlier findings (**Figure 1B**), confirming the robustness and validity of our classification approach.

To correlate metabolic dynamics with *L. pneumophila* replication in infected macrophages (**Figure 1C**), we chose to measure various mitochondrial parameters as we had previously shown that *L. pneumophila* manipulate mitochondrial functions to its advantage (23, 40). Our miniaturized imaging approach detected the limits of each single, living cell using the blue channel and tracked bacterial replication dynamics of the Lpp-WT and -Δ*dotA* strains using the green channel as described above (**Figure 1A**). The two remaining free fluorescence channels (red and far-red) were used to measure metabolic outputs in a multiplex approach (**Figure 2A**). Altogether we measured in each cell the mitochondrial mass, the mitochondrial morphology, the Δψm, early and late cell death, and ROS levels using dyes for living cells (**Table 1**). Furthermore, we measured morphological parameters such as nuclear and cellular size and roundness. BATLI was then used to reconstruct bacterial replication dynamics and to classify macrophages at the end of the infection cycle into the two previously defined categories: those supporting bacterial replication (Class 1) and those not (Class 0) (**Figure 1C** and **2B**). The progression in time of mitochondrial and cellular parameters for each classified cell was blotted using BATLI, as graphs that were color-grouped according to their specific bacterial-replication class (**Figure 2B, 2C, 2D**, and **S2A-I**). This allowed us to track in every infected cell the progression of cellular parameters and to correlate them with bacterial replication outcomes.

**Figure 2.**
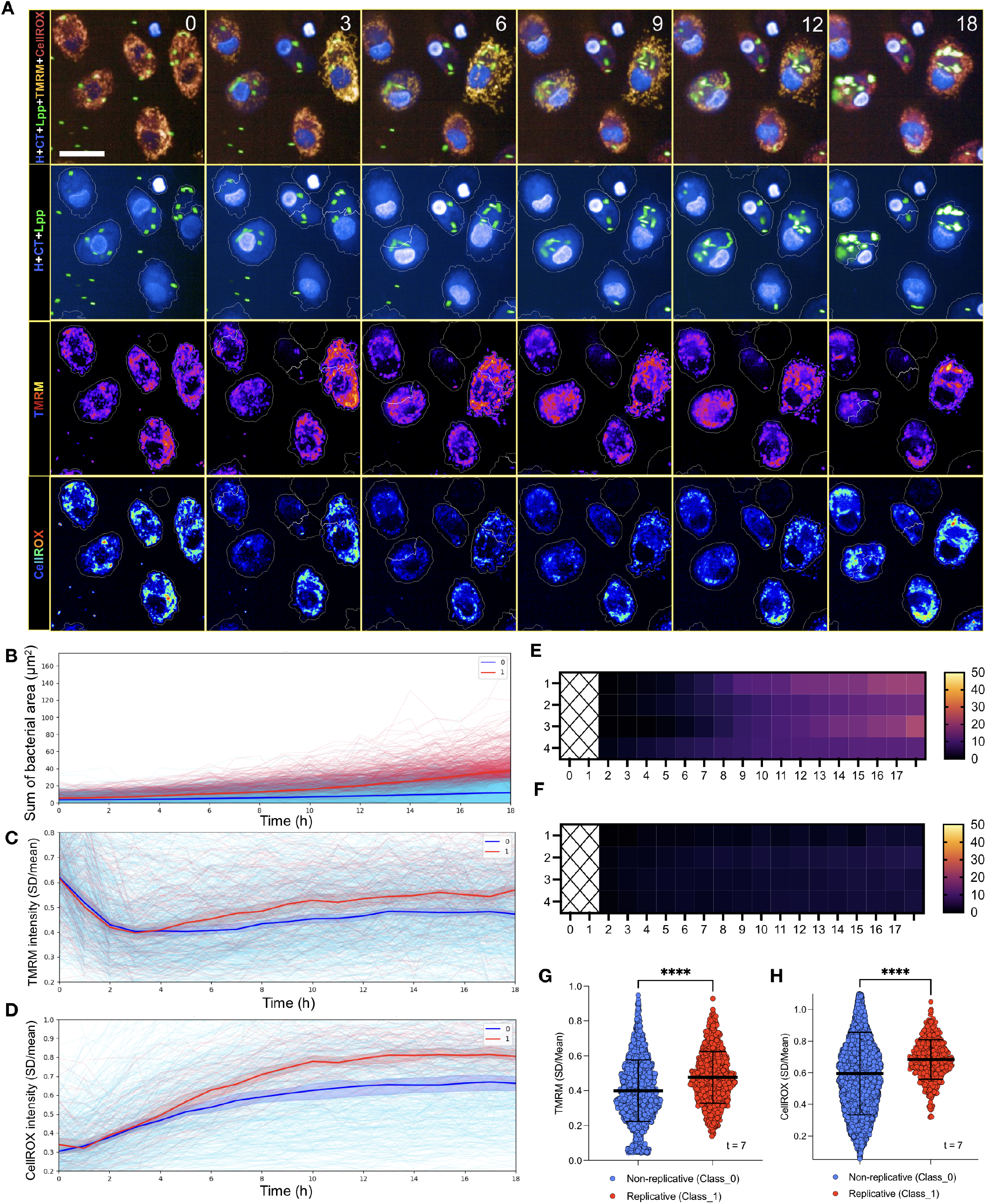
Single-cell trajectories reveal that early alterations of mitochondrial functions occur preferentially in macrophages supporting *L. pneumophila* replication before bacterial replication starts. (**A**) Confocal images of infected hMDMs during the infection course, from 0 to 18 hours hpi. GFP-expressing Lpp are shown in green. In blue are shown the nuclei stained with Hoechst (H) and cytoplasm with Cell Tracker Blue (CT). The red channel was used for TMRM and the far-red channel for CellROX. Smaller boxes in the bottom contained separated channels. Here, TMRM and CellROX are shown as pseudo color palettes to highlight intensity features. Bar = 20 µm. (**B**) Single-cell tracks of bacterial vacuole sizes (µm2) for Lpp WT-infected hMDMs (0 to 18 hpi). Each single line represents one cell. Single-cell trajectories were classified using BATLI as hMDMs with replicative vacuoles (shown in red) and hMDMs with non-replicative vacuoles (blue). (**C**) Backtracking analysis of Δψm measurements using TMRM signal from time lapse images in BATLI-classified Lpp WT-infected single-cell trajectories (infected hMDMs, 0 to 18 hpi). Class 1 (red) = infected hMDMs showing replication of bacteria, Class 0 (blue) = infected hMDMs showing no replication of bacteria. (**D**) Same than (C) but analyzing cellular ROS measurements using CellROX signal. (**E**) Heat map showing the percentage of hMDMs where Lpp-WT bacterial area doubled with respect to the mean area of previous time points (minimum 2 consecutive time points). Each row is a dataset of an independent infection experiment. Legend values are % of total cells (**F**) Same than E but for hMDMs infected with avirulent Lpp-Δ*dotA* mutant strain. (**G**) TMRM signal (SD/Mean) of classified hMDMs in B at 7 hpi (****= p<0.0001, unpaired t test). (**H**) CellROX signal (SD/Mean) of classified hMDMs in B at 7 hpi (****= p<0.0001, unpaired t test).

Two cellular parameters in hMDMs were identified as correlated with bacterial replication. One was Δψm, which was measured as the aggregation of Tetramethylrhodamine, methyl ester (TMRM) dye into mitochondria (SD/Mean (47)). TMRM is a cell-permeant dye that is sequestered at mitochondria depending on the Δψm. The analysis of the Δψm over time showed that after an initial reduction in all Lpp-WT-GFP-infected cells, permissive macrophages showed higher and sustained levels of Δψm over time (**Figure 2C**). The second parameter we identified was ROS. We observed that infected macrophages with virulent or T4SS-defective *L. pneumophila*, showed an overall increase in the aggregation of CellROX^®^ dye in intracellular organelles (SD/Mean). CellROX^®^ is a fluorogenic probe for measuring oxidative stress in living cells. However, those containing replicative vacuoles had higher levels of CellROX^®^ signal compared to macrophages where no bacterial replication was observed (**Figure 2D**). This suggests that cellular ROS are differentially regulated in infected cells that support and do not support *L. pneumophila* replication. The remaining parameters we had measured, such as mitochondrial morphology (**Figure S2A** and **S2B**), mitochondrial mass (**Figure S2C**), cell size (**Figure S2D**), nuclei size (**Figure S2E**), nuclei condensation (**Figure S2F**), nuclei morphology (**Figure S2G**), or cell morphology (**Figure S2H**) were not different between macrophages supporting bacterial replication (Class 1) or not (Class 0). Thus, sustained Δψm and elevated ROS are unique cellular markers correlated with the ability of infected macrophages to facilitate intracellular replication of *L. pneumophila*.

As we aimed to predict whether a given human macrophage will support bacterial replication or not before bacterial replication actually starts, we needed to determine the time points when intracellular bacterial replication of *L. pneumophila* in hMDMs starts in most of the cells. For this, we analyzed single-cell trajectories of hMDMs infected by the Lpp-WT strain (**Figure 1D** and **1E**) and calculated the percentage of macrophages where the bacterial area at least doubled with respect to the mean area at previous time points. We observed that no significant changes occurred until 8 hpi (**Figure 2E** and **Table S1**). Furthermore, same calculation using Lpp-Δ*dotA*-infected macrophages showed no duplication of bacteria across the whole time-course (**Figure 2F**), validating the approach. We thus chose 7 hpi as the time point when bacterial replication starts. Taking this definition, we revealed that differences between both classes of macrophages (allowing bacterial replication or not) were indeed already significantly different at 7 hpi for Δψm and ROS (p<0.00005, **Figure 2E** and **2F**).

To understand where the measured ROS was produced we visually inspected the images obtained (**Figure 2A**) what suggested that the ROS measured in the infected macrophages might be produced at mitochondria (mROS). To follow up this observation we stained hMDMs simultaneously with Mitotracker^™^ Deep Red (a fluorescent probe used to visualize mitochondria) and CellROX^®^ Orange (to measure ROS production, **Figure 3A**). Spatial correlation analyses of fluorescence signals in living cells showed similar profiles for Mitotracker^™^ Deep Red and CellROX^®^ Orange, and clearly different profiles compared to Hoechst signal (**Figure 3B** and **3C**). To further investigate the localization of ROS production, we conducted assays with drugs targeting the mitochondrial Electron Transport Chain (ETC). Specifically, we treated hMDMs with Rotenone and Antimycin A, which are inhibitors of complex I and complex III of the mitochondrial ETC, respectively, because these are the main sites where mROS is produced (48). We then simultaneously measured Δψm and ROS production at the single cell level (**Figure 3D-G**). By inhibiting ETC complex I or complex III, we disrupted the flow of electrons that passes through each of these complexes at the ETC. As a result, electrons accumulate at the site of inhibition and subsequently leak partially, reducing O_2_ and generating superoxide radicals O_2_·in mitochondria, *i*.*e*. mROS. As control we measured ROS by CellROX^®^ dye in non-infected (NI) hMDM in a time course experiment showing that ROS levels increased slightly without infection (**Figure 3D**). Treatment of cells with Antimycin A did not change ROS levels compared to non-treated cells (**Figure 3E**), while cells treated with Rotenone or a combination of Antimycin A and Rotenone showed a two-fold increase of ROS levels upon drug addition (**Figure 3F** and **3G**). As we observed a direct effect on ROS production upon inhibition of ETC complex I with Rotenone, our experiments strongly suggested that measuring ROS using CellROX^®^ dye in hMDMs is measuring mROS production by complex I in the mitochondria, but not by complex III.

**Figure 3.**
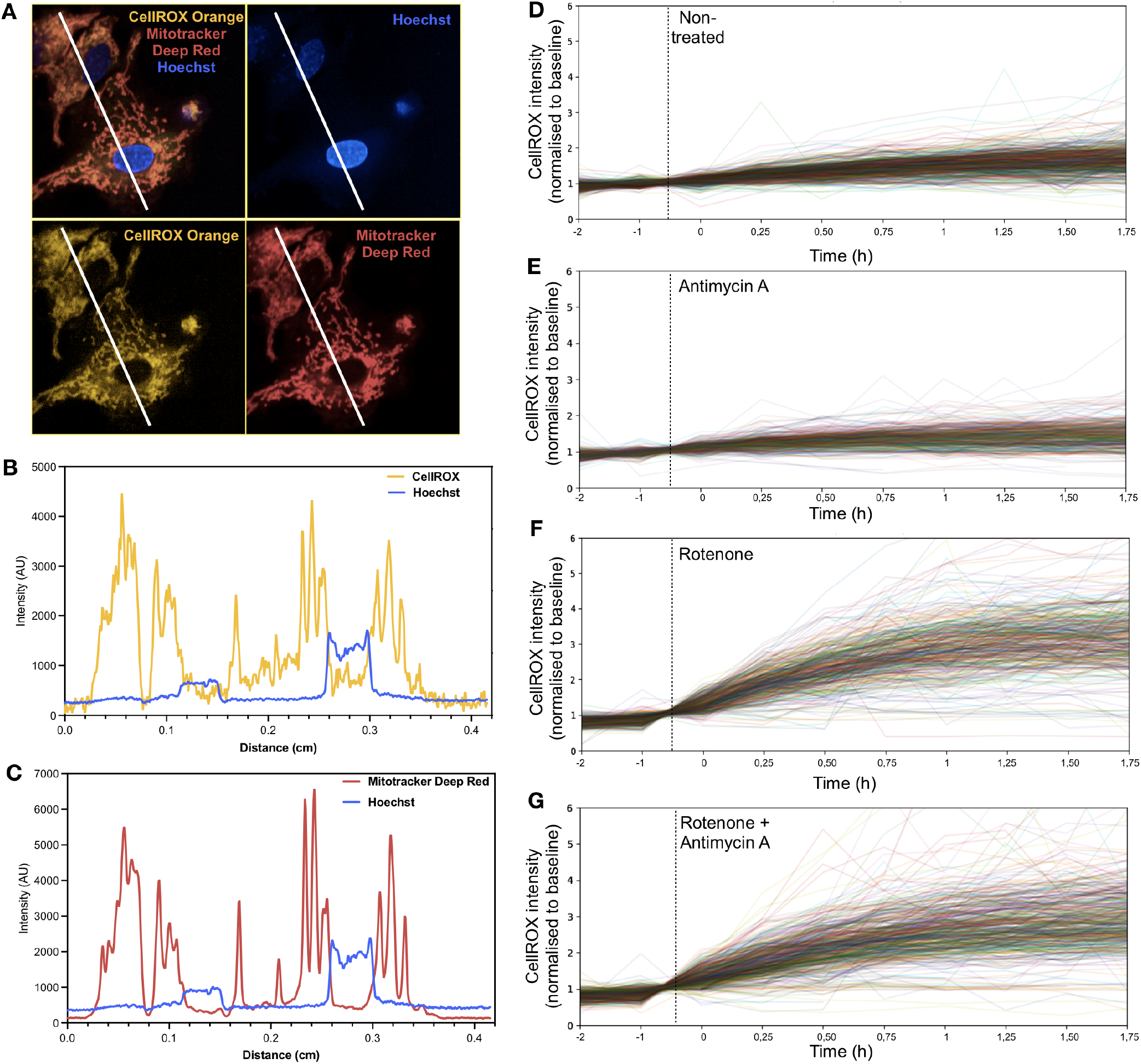
ROS measurements in hMDMs are mROS produced by mitochondrial electron transport chain (ETC) complex I, but not by complex III. (**A**) Confocal images of hMDMs stained with CellROX Orange to measure cellular ROS (yellow), Mitotracker Deep Red to stain mitochondria (red) and Hoechst (nuclei). Spatial correlation analyses of fluorescence signals were carried out on the white line. (**B**) Intensity profiles from spatial correlation analyses for CellROX Orange and Hoechst. (**C**) Same than (B) for Mitotracker Deep Red and Hoechst. (**D**) Single-cell trajectories of ROS measurements (CellROX) in non-treated hMDMs. Dotted line shows the time when medium (control) was added. Values are normalized to the baseline (−2 to 0 h). (**E**) Same than (D) in Antimycin-A-treated hMDMs. Dotted line shows the time when treatment was added. (**F**) Same than (E) in Rotenone-treated hMDMs. (G) Same than (E) in Rotenone+Antimycin-A-treated hMDMs.

Taken together, cells within the same replication class (Class 1/0) exhibited a similar evolution of several cellular parameters throughout the course of infection. However, sustained Δψm and elevated mROS are key metabolic markers that correlate with the ability of infected macrophages to facilitate bacterial intracellular replication.

### *Infection with a Type IV secretion system deficient* L. pneumophila *strain increases resistance of macrophages to induced cell death*

Another parameter highly dependent on mitochondria during infection is the induction of cell death, therefore we used BATLI to study macrophage cell death during infection by comparing replication competent (Lpp-WT) and replication deficient (Δ*dotA*) *L. pneumophila* strains. hMDMs were infected with Lpp-WT-GFP and Lpp-Δ*dotA*GFP strains and stained with Annexin-V or SITOX as a markers of early and late cell death (49, 50), respectively. Images were taken hourly, up until 18 hpi. Following image capture, we used BATLI to classify infected macrophages according to bacterial replication, as described above (**Figure 1E** and **2B**). While the dynamics of late cell death were not different between both classes of macrophages (**Figure S2I**), judged by the lack of differences in nuclei condensation (**Figure S2F**), early cell death preferentially happened in those infected cells where bacteria did not replicate (**Figure 4A**, Class 0, blue lines). In contrast macrophages supporting bacterial replication highly delayed the onset of cell death events (**Figure 4A**, Class 1, red lines). To follow this result, we used BATLI this time to classify infected macrophages according to early-celldeath-fate (**Figure 4B**). Thus, in these analyses, each single cell trajectory was classified as triggering early cell death (Class 1, increased Annexin-V signal over threshold) or not (Class 0, non-increased Annexin-V signal). In 75% of LppWT-GFP-infected macrophages, the intensity of Annexin-V increased over time, and these were thus classified as Class 1 (**Figure 4B**). Moreover, cell death was preceded by a reduction in Δψm specifically in those infected cells that then died (**Figure 4C**). Thus, early cell death is a common fate occurring during the course of infection specifically in those cells where bacterial replication is not supported.

**Figure 4.**
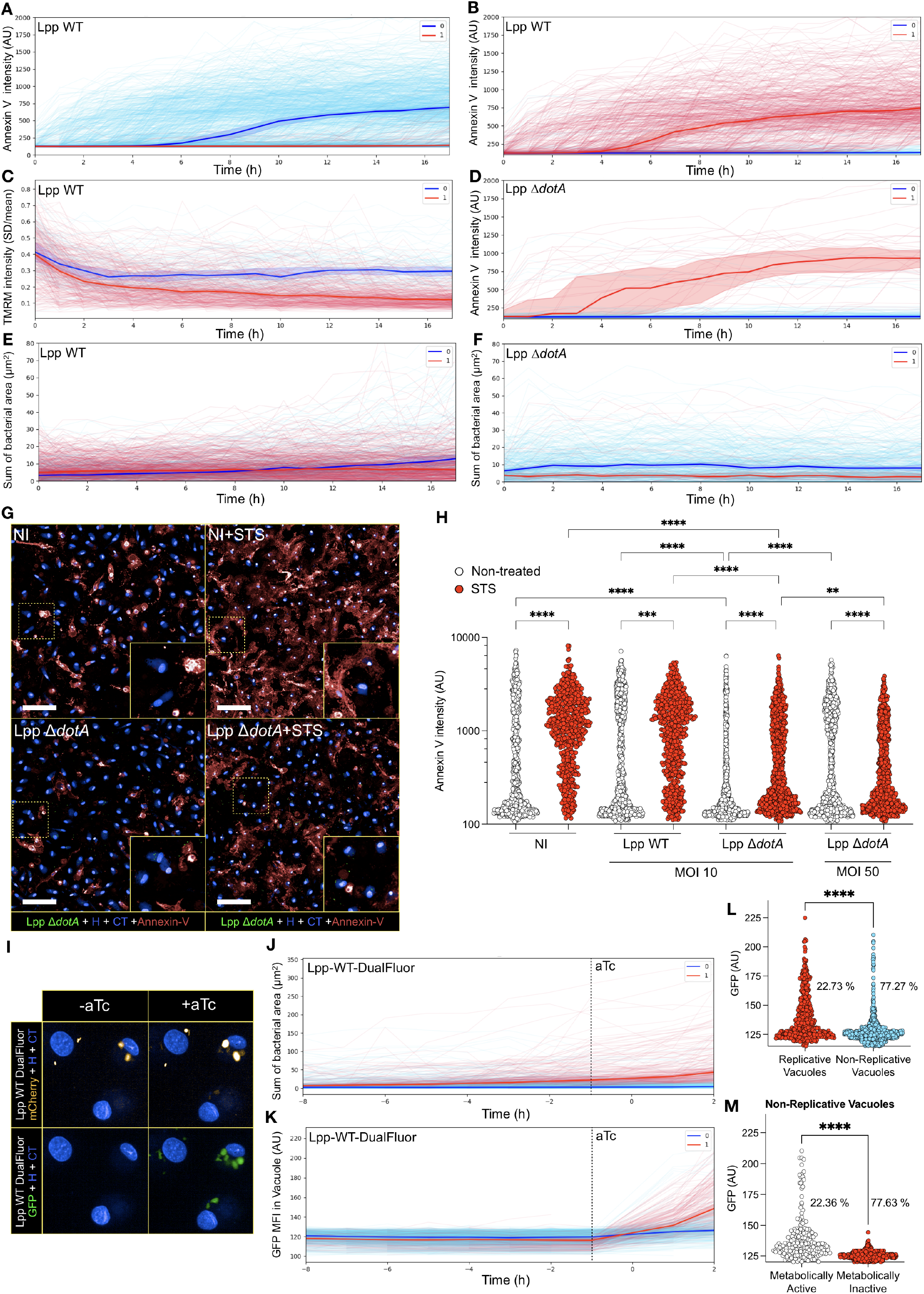
Lpp-Δ*dotA* protects infected cells from exogenously triggered cell death and non-replicative Lpp-WT vacuoles still have metabolically active bacteria. (**A**) Backtracking analysis of early cell death measurements using Annexin-V signal from time lapse images in BATLI-classified Lpp WT-infected single-cell trajectories. Class 1 (red) = infected hMDMs showing replication of bacteria, Class 0 (blue) = infected hMDMs showing no replication of bacteria. (**B**) Classification of single-cell trajectories of Lpp WT-infected cells according to early cell death measurements (Annexin V signal) using BATLI. Class 1 (red) = infected hMDMs showing early cell death, Class 0 (blue) = infected hMDMs not showing early cell death. (**C, D**) Same than (B) but in Lpp-Δ*dotA*-infected hMDMs. (**E**) BATLI analysis of bacterial area (vacuole sizes in µm^2^) upon classification of single-cell trajectories according to early cell death measurements (B) in Lpp WT-infected hMDMs. Class 1 (red) = infected hMDMs showing early cell death, Class 0 (blue) = infected hMDMs not showing early cell death. (**F**) Same than (E) in Lpp-Δ*dotA*-infected hMDMs. Class 1 (red) = infected hMDMs showing early cell death, Class 0 (blue) = infected hMDMs not showing early cell death. (**G**) Confocal images of infected hMDMs. GFP-expressing Lpp are shown in green. In blue are shown the nuclei stained with Hoechst (H) and cytoplasm with Cell Tracker Blue (CT). Annexin-V staining is shown in red. Cells were infected with Lpp-WT or Lpp-Δ*dotA* at MOI 10, with Lpp-Δ*dotA* at MOI 50, or kept non-infected (NI). At 5 hpi, Staurosporine (1µM) was added or not. Images at 10 hpi are shown. Bar = 100 µm. (**H**) Cell death was determined at 10 hpi by measuring Annexin 647 intensity (AU) in all conditions (*= p<0.05, **=p<0.01, ***=p<0.001, ****= p<0.0001, ordinary one-way ANOVA). (**I**) Confocal images of hMDMs infected with Lpp-WT-DualFluor strain. This strain constitutively expresses mCherry fluorescent protein and has the expression of GFP under the control of a Anhydrotetracycline (aTc)-inducible promotor. At 16 hpi, aTc was added to reveal the metabolic status of intracellular bacteria, as only metabolically active bacteria would express GFP. (**J**) BATLI classification of bacterial vacuoles according to vacuole growth. Class 1 = replicative vacuoles, Class 0 = non-replicative vacuoles. (**K**) Backtracking of GFP mean fluorescent intensity in the vacuoles classified in (F). (**L**) Measurements of GFP signal (AU) and percentages of each population of vacuoles classified in F (****= p<0.0001, unpaired t test). (**M**) Measurements of GFP signal (AU) and percentages of each population of non-replicative vacuoles according to the increase of GFP signal (****= p<0.0001, unpaired t test).

In contrast, early cell death was only observed in 10% of Lpp-Δ*dotA*-GFP-infected cells, confirming that early cell-death is T4SS-dependent as previously suggested (51) (**Figure 4D**). In these WTand Δ*dotA*-infected cells classified according to their fate in cell death, we tracked back the GFP area as a proxy of *L. pneumophila* replication. This revealed that the bacterial area occupied by Lpp-WT was similar in the two classes of macrophages at the beginning of the infection (0 to 8 h), suggesting that initial differences in bacterial uptake did not correlate with specific cell-death fates in infected macrophages (**Figure 4E**). The bacterial area of Lpp-WT increased at late times post-infection (8 to 18 h) in those cells not triggering cell death (Class 0), as previously suggested (40). Thus, bacterial replication (i.e. increment of intracellular bacterial area) occurred specifically in those WT-infected hMDMs not triggering cell death. During infection with the Lpp-Δ*dotA* mutant, the bacterial area did not increase over time in either of the two distinct celldeath-fate classes, as expected. Surprisingly, those cells that died had a lower bacterial load than the infected cells avoiding cell death (**Figure 4F**). This was intriguing as one would expect that higher bacterial loads, would increase cell death in infected cells. However, strikingly, our result suggested that higher bacterial loads of the avirulent Lpp-Δ*dotA* strain reduced cell death of infected macrophages. To investigate this surprising finding, we infected hMDMs with Lpp-WT or Lpp-Δ*dotA* at different multiplicities of infection (10 and 50 MOI), or left them uninfected (non-infected, NI). At 5 hpi we treated or not the cells with 1µM Staurosporine (STS), a well-known inducer of cell death (52), and measured Annexin-V intensity at 10 hpi (**Figure 4G**). As shown in **Figure 4H**, the percentage of single cells showing high Annexin-V intensity upon STS treatment of noninfected or WT-infected cells, was significantly higher compared to the percentage of cells showing high Annexin-V in Lpp-Δ*dotA*-infected hMDMs (NI+STS vs. Lpp-Δ*dotA*+STS p < 0.0001, Lpp-WT+STS vs. Lpp-Δ*dotA*+STS p < 0.0001). This suggested that infection with the Lpp-Δ*dotA* strain protected from STS-induced cell death. Moreover, infection of hMDMs with Lpp-Δ*dotA* at higher MOI reduced the acquisition of the Annexin-V marker even more (Lpp-Δ*dotA* MOI 10 vs. Lpp-Δ*dotA* MOI 50, p < 0.0001), suggesting that high infections with a T4SS-deficient mutant strain protected hMDMs against STS-induced cell death.

In summary, our results suggest that resistance to cell death from exogenous stimuli might be triggered when macrophages are infected with a *L. pneumophila* strain having a non-functional T4SS. These results need further analyses, but they highlight the insights one can obtain by analyzing single cell dynamics using BATLI in the context of host-pathogen interactions.

### *Backtracking of intracellular bacterial vacuoles reveals that heterogeneity in the metabolic activity of* L. pneumophila *also impacts infection*

It can be argued that the heterogeneity in macrophages analyzed with BATLI, where intracellular replication of *L. pneumophila* occurs in only a portion of all infected cells (**Figure 1B**), might also be influenced by the metabolic status of the bacteria themselves. Therefore, we used BATLI to also analyze the metabolic heterogeneity of the bacteria during infection and to understand whether the metabolic status of the pathogen has an impact on their replication capacity (**Figure 1B** and **1E**). We constructed a *L. pneumophila* strain (Lpp-DualFluor) constitutively expressing mCherry fluorescent protein and the expression of GFP under the control of an Anhydrotetracycline (aTc)-inducible promotor. We infected hMDMs with the Lpp-DualFluor strain and followed intracellular replication of *L. pneumophila* using the mCherry signal. At a late time point of the infection course (16 h), we added aTc to reveal the metabolic status of the intracellular bacteria, as only metabolically active bacteria would transcribe and translate GFP (**Figure 4I**). Using BATLI, we tracked bacterial vacuoles in the infected cells and classified the bacterial vacuoles according to vacuolar growth (**Figure 4J**). Class 1 (replicative vacuoles) and Class 0 (non-replicative vacuoles) were defined as in **Figure 1E** and **2B**, but this time using mCherry signal to segment intracellular bacterial vacuoles. Then we backtracked GFP mean fluorescent intensity in the vacuole, which corresponds to the activity of the aTc-inducible promotor, in these two classes (**Figure 4K**). This analysis matched the levels of metabolic activity of bacteria with their corresponding replicative status. Then we used BATLI to classify bacterial vacuoles according to the increase of GFP signal growth upon adding aTc inducer (**Figure S3A**) and backtracked the dynamics of the bacterial vacuole area (**Figure S3B**). We then merged the two datasets and reconstructed the trajectories by matching every single vacuole at every time-point using its unique ID, what allowed us to determine that 23% of vacuoles were replicative (Class 1) and 77% were non-replicative (Class 0), according to the significantly different GFP signal (139 ± 20 AU vs. 127 ± 12 AU, respectively, p < 0.0001, **Figure 4L**). Interestingly, the analysis of the population of non-replicative vacuoles showed that 22% increased GFP upon aTc addition (metabolically active bacteria), while 78% did not (metabolically inactive), having also a GFP signal significantly different between them as expected (138 ± 20 AU vs. 123 ± 5 AU, respectively, **Figure 4M**). These results indicated that most of the nonreplicative vacuoles contain metabolically inactive bacteria, that probably had been killed by the macrophage. However, a small percentage of non-replicative vacuoles (22%) harbored metabolically active bacteria although they did not replicate, suggesting that these might contain dormant bacteria. Whether the metabolic dynamics of infected macrophages might induce bacterial dormancy is a question that deserves further investigation. In summary, our results suggest that the metabolic status of the pathogen also impacts the heterogeneity observed in the replication rate of *L. pneumophila* in macrophages (**Figure 1B** and **1E**), showing that our approach using BATLI is a powerful tool to disentangle the contributions of each player to the outcome of an intracellular infection.

### Using backtracking datasets to train a machine-learning model allows the prediction of whether macrophages will support bacterial replication before the onset of replication

By using BATLI, we were able to classify cells depending on the infection outcomes, i.e. replication of bacteria or not, and directly link the resulting classes with cell dynamics prior to the onset of bacterial replication. The data generated with BATLI (example in **Table S2**) can be easily used to further train machine-learning models to investigate new questions.

Our next aim was to predict replication of Lpp-WT in hMDMs, before the onset of bacterial replication (7 hpi, **Figure 2E, 2F** and **Table S1**), by using our single-cell datasets of tracked bacterial infection provided by BATLI. For this we used Δψm and mROS, as these metabolic parameters were significantly different before the onset of replication (7 hpi, **Figure 2G** and **2H**). We multiplexed them in living assays, and trained a machine-learning model using these parameters. To construct the model we used the logistic regression method, a supervised machine learning algorithm widely used for binary classification (53). We pooled the datasets from three independent infection experiments and trained the model with the trajectories of 5,119 Lpp-WT-infected macrophages (training dataset). Then, we used the model to predict the fate of 1,707 trajectories of Lpp-WT-infected macrophages that the model never saw before (validation dataset). We trained the model using different trajectory lengths, resulting that the best predictive capacity of the validation dataset was achieved when the model considered single-cell trajectories between 0 and 5 hpi (**Figure 5A** and **S4**). In this validation dataset, 83% of true permissive macrophages were classified by the trained model as such, and 60% of the true repressive macrophages were classified as such, as can be seen in the confusion matrix depicted in **Figure 5B**. As an explainable machine-learning model, the probability of any single, infected cell of being supportive of *L. pneumophila* replication can be determined by the logistic equation and Δψm and mROS SD/mean values (**Figure 5C**). The overall performance of the model is summarized by its AUROC (area under ROC curve) as well as its precision-recall curve (**Figure 5D** and **5E**, respectively). An AUROC value close to 1 suggests an ideal model, while values around 0.5 indicate a random model. Our model achieved an intermediate value of 0.74, which is a good score for prediction performance.

**Figure 5.**
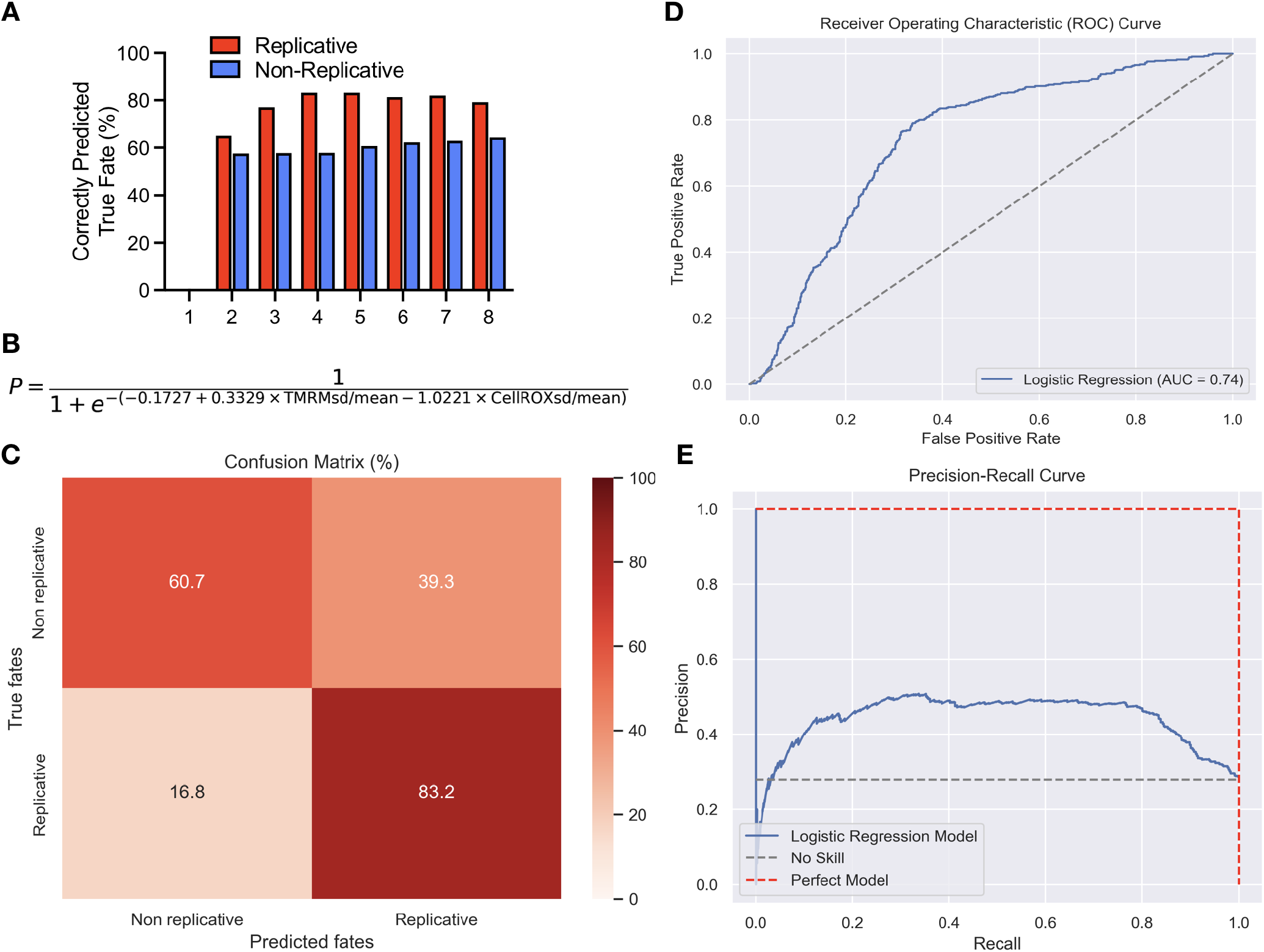
BATLI analyses combined with machine learning allows the prediction of replication outcomes in single-cells before the on-set of replication. (**A**) Predictive accuracy across different trajectory lengths. The bar plot shows the percentage of correctly predicted true fates (y-axis) for both replicative (red) and non-replicative (blue) macrophages across different trajectory lengths (x-axis, hpi). (**B**) Logistic regression equation describing the probability (P) of a single infected macrophage supporting *L. pneumophila* replication. This probability is derived from Δψm (TMRM SD/mean) and mROS (CellROX SD/mean) values. (**C**) Receiver Operating Characteristic (ROC) curve for the random forest model predicting replicative vacuoles in *L. pneumophila*-infected macrophages. The X-axis represents the false positive rate (1-specificity), while the Y-axis represents the true positive rate (sensitivity). AUC = Area Under Curve. (**D**) Precision-recall curve showing model performance, with the logistic regression model (blue curve) compared to a no-skill model (gray dashed line) and a hypothetical perfect model (red dashed line).

In summary, the datasets obtained and analyzed using BATLI allowed to successfully train an explainable machinelearning model using the logistic regression algorithm and the mitochondrial parameters Δψm and mROS. This model predicted with 83% accuracy, at the single-cell scale and in living cells, which infected human macrophages will be permissive to *L. pneumophila* replication at time-points where bacterial replication has not yet started (5 hpi), confirming that the here identified parameters, Δψm and mROS, are excellent predictable markers for cells supporting bacterial replication.

## Discussion

One fundamental question in infection biology we explored is: *Can we predict infection outcomes at the single cell level?* By facilitating a detailed analysis of cellular dynamics and outcomes, our unique approach allowed us to investigate into the concept of computational reducibility within hostpathogen interactions. Computational reducibility of a system is a concept introduced by Stephen Wolfram (54) that refers to the idea that, if the system is computationally reducible, the outcome of this complex system can be predicted in a simpler or shorter way than simply observing its natural evolution. Our results indicated that bacterial infection exhibits pockets of computational reducibility at the single-cell level, potentially opening new avenues to predict the progression of infectious diseases. This finding suggests that certain aspects of the complex dynamics in host-pathogen interactions are indeed predictable, moving beyond traditional infection biology approaches and shedding new light on how predictive biology can decipher these complexities.

Previous efforts to predict transitions in single-cell states during infection have predominantly used inference methods on transcriptomic datasets, particularly single-cell RNA sequencing (sc-RNA-seq) and metabolomic datasets, to elucidate the metabolic shifts occurring within infected cells (55– 58). Others focused their research on the variability within bacterial populations during infection, seeking to understand how these differences influence the infection process (59, 60). Despite the advancements provided by sc-RNA-seq and metabolomics in revealing metabolic dynamics of hosts and pathogens, these technologies cannot capture the dynamics of inherently heterogeneous processes like infection because metabolic patterns vary and evolve in individual infected cells over time.

To construct a model allowing to predict transitions in single-cell states and to study macrophage heterogeneity during bacterial infection, we combined high-throughput confocal image analyses and dyes for living cells with the development of a new software, named BATLI, to reconstruct the trajectories of metabolic parameters in single, infected cells. Indeed, several intracellular bacteria, despite the inherent defense mechanisms of macrophages, can infect and proliferate within these immune cells thereby causing human disease (61). Here we developed an approach to perform retrospective analyses of infected human primary macrophages to associate specific cellular fates to earlier cellular conditions, at the single cell scale. To investigate single-cell fates during infection of macrophages we used *L. pneumophila*, the intracellular bacterium that is the causative agent of Legionnaires’ disease (16) as a model. BATLI, a tool we developed here, was used to classify infected human primary macrophages according to cellular fates (bacterial replication or not, cell death or not) that are only known at the end of the infection, and then to track back every infected cell to analyze earlier metabolic parameters unambiguously linked to these cell fates.

Our investigation led to intriguing observations. First, *L. pneumophila* replicates within some human primary macrophages but not in others, pointing towards the heterogeneity of macrophages responses as a potential explanation for this observation. Indeed, bacterial replication only occurs in 20% of *L. pneumophila*-infected macrophages, indicating that it is the exponential growth of the pathogen within this subset of permissive, infected cells that allows *L. pneumophila* to overcome the success of macrophages in killing intracellular bacteria.

Our findings revealed that the maintenance of Δψm during infection, as well as the delay of cell death, unambiguously and preferentially occurred in those macrophages that supported bacterial replication. This result corroborated our previous results that human macrophages infected with *L. pneumophila* WT sustain the Δψm in conditions where OXPHOS was reduced (40). This phenomenon was found to be T4SS dependent, and the bacterial effector LpSpl was shown to be partially involved in the induction of the ‘reverse mode’ of the mitochondrial ATPase, which sustained the Δψm levels and delayed cell death in infected cells (40). The use of BATLI extended this understanding by demonstrating that among cells infected by *L. pneumophila* WT, only those cells in which bacterial replication occurs at the end of the infection (permissive macrophages) retained their Δψm since the beginning of the infection, which was combined with an increased production of mROS. The increase in mROS corroborated the known metabolic reprogramming of macrophages in response to Pathogen Associated Molecular Patterns (PAMPs). Under these conditions, mROS production is involved in cytokine production and antimicrobial activities to eliminate infection. Typically, mROS accumulation might trigger inflammasome activation, which ultimately leads to pyroptosis and cell death to prevent bacterial spread (10). However, our findings indicate that cell death is delayed in cells harboring replicative vacuoles despite the increase of mROS, which might indicate that bacterial-induced maintenance of Δψm circumvent the induction of cell death triggered by mROS production. Moreover, these results suggest that the functioning of mitochondrial ETC Complex I is affected during Lpp-WT infection, which leads to the production of mROS, and only those macrophages showing this specific response have a metabolic status compatible with the intracellular replication of *L. pneumophila*. Thus, the diversity in mitochondrial parameters between cells plays a significant role in determining the success or failure of an infection at the single-cell level.

Additional findings arising from our analyses suggest that a program increasing resistance to exogenously induced cell death exists in human macrophages, and this program can be triggered by infecting macrophages with an avirulent but life *L. pneumophila* strain, the Δ*dotA* mutant, which possesses all PAMPs but does not inject T4SS effectors. It was previously shown that Staphylococcus aureus-infected hMDMs have a higher resistance to STS-induced cell death compared to their uninfected counterparts (62), which is caused by the up-regulation of the anti-apoptotic factor Mcl-1. Whether the protective effect of Lpp-dotA-infection involves similar antiapoptotic pathways is an intriguing question that deserves further investigation.

Our approach also allowed us to investigate the impact of the metabolic status of the pathogen on the replicative status of the vacuoles, showing that most of non-replicative vacuoles are populated by metabolically inactive bacteria, and only a minor portion of them seem to be dormant bacteria. These results highlight the existence of a minor population of metabolically active and non-replicative *L. pneumophila* inside human macrophages that might be those that had been defined as dormant or persistence bacteria (63). BATLI is a tool that can be used to study this phenomenon, which opens the possibility of studying at the same time the metabolic conditions of host cells and bacteria and explore interesting questions such as whether specific metabolic conditions of infected cells induce dormancy or persistence in the population of intracellular bacteria. Altogether, our results propose a model where metabolic dynamics of infected cells critically dictate whether intracellular pathogens will succeed in causing fruitful infections and seems excellent to study the dynamics of cell-fate decisions at the single cell level to disentangle the contribution of each player, host cell and pathogen, in the final outcome of an infected macrophage. However, our study extends significantly beyond retrospective analysis of metabolic parameters as our robust data-gathering and cell-fate-classification approach provides highly curated and labeled datasets, which are invaluable for training machinelearning and prediction models. Indeed, we generated dynamic datasets that are essential for testing hypotheses within the framework of predictive biology.

A previous study also leveraged automated high-content imaging to predict infection outcomes. Voznica *et al*. conducted an investigation into the susceptibility of epithelial cells to *Salmonella* Typhimurium infection (64). Their approach involved developing a mathematical model to predict the likelihood of a single, infected cell of being re-infected. The model achieved a prediction accuracy of 62% for HeLa cells and 66% for Caco-2 cells in terms of vulnerability to subsequent *Salmonella* infections. In our study, we advanced this predictive framework by utilizing BATLI in combination with logistic regression algorithms to predict bacterial replication within *L. pneumophila*-infected human primary macrophages. Our model demonstrated a significantly higher prediction accuracy of 83% for determining whether an infected macrophage will support bacterial replication at a time-point, 5h post-infection, when *L. pneumophila* intracellular replication have not started yet. The high accuracy of our model might indicate the importance of integrating comprehensive single-cell trajectory data with advanced computational techniques using explainable, shallow learning algorithms such as logistic regression. Our model achieved an AUROC value of approximately 0.74. While this value is significantly less than 1, indicating that Δψm and ROS —the two features we identified as determinants of bacterial replication— are not sufficient to fully predict the fate of an infected cell, they still contribute substantially to prediction accuracy. It is likely that other metabolic parameters also influence this fate, as can be expected in such a complex system. Nonetheless, Δψm and ROS alone achieve a high level of predictive accuracy, highlighting their prominent role in determining bacterial replication fate. The ability of highcontent imaging to capture dynamic and quantitative measurements of cellular parameters over time, along with the robust classification and backtracking capabilities developed for this study, might be pivotal for developing more precise predictive models in infection biology.

BATLI might also have important utility beyond the field of infection biology. It could be applied to other research fields such as cancer biology or developmental biology. In cancer research, for example, our approach could be used to study how early cellular parameters predict specific cellular fates, such as cell division or transformation. Similarly, in developmental biology, our tools could help to elucidate how certain cellular dynamics correlate with developmental processes at the single-cell level.

In summary, our study demonstrates the power of combining high-throughput live-cell imaging and computational tools to elucidate the metabolic dynamics governing infection outcomes at the single-cell level. By leveraging BATLI and machine learning, we successfully predicted infection fates with high accuracy, highlighting early mitochondrial alterations as key predictive markers and providing detailed insights into the cellular dynamics involved in the replication of intracellular bacteria within human macrophages.

## Acknowledgements

We acknowledge C.B.’s lab members for fruitful discussions. We thank Stéphane Rigaud (Image Analysis Hub) for guidance with programming and data analysis. We thank Nathalie Aulner, Anne Danckaert, Nassim Mahtal and the Photonic BioImaging (PBI) UTechS at Institut Pasteur for support. We acknowledge Francisco Garcia del Portillo (CNB-CSIC, Madrid, Spain) and Francisco Rodriguez Garcia (Institut Pasteur) for the critical reading of the manuscript. We also acknowledge Hélène Laude and the ICAReB team for their support providing human blood samples. This research was funded by the Institut Pasteur, the Agence National de Recherche (grant number ANR-10LABX-62-IBEID to C.B. and ANR21-CE15-0038-01 to P.E.), the Programmes Transversaux de Recherche (PTR) (grant number PTR-651) from Institut Pasteur to P.E, the Fondation de la Recherche Médicale (FRM) (grant number EQU201903007847) to C.B., and the Région Ile-de-France (program DIM1Health) to PBI (part of FranceBioImaging, ANR-10-INSB-04–01). J.E.M. was supported by PasteurRoux-Cantarini Postdoctoral Fellowship. M.D. was supported by the Ecole Doctorale FIRE – ‘Programme Bettencourt’. Figures contain elements from BioRender.

## Methods

### Cell culture of human monocytes derived macrophages (hMDMs)

Blood from healthy donors was provided by either the French National Blood Service (Etablissement Français du Sang – EFS) or by the Clinical Investigation and Access to Biological Resources Service (Investigation Clinique et Accès aux Ressources Biologiques – ICAReB) at Institut Pasteur Paris. Peripheral blood mononuclear cells (PBMCs) were isolated from the rest of the blood components by Ficoll density gradient centrifugation (Lymphocyte Separation Medium Eurobio Scientific) at room temperature. Monocytes were positively sorted using CD14 magnetic beads (CD14 MicroBeads Miltenyi Biotec). Following, monocytes were differentiated into hMDMs by culturing them in X-VIVO15 medium without Phenol Red (Lonza) and with recombinant human M-CSF (rh-MCSF Miltenyi Biotec) at a concentration of 25 ng/ml for 6 days at 37°C with 5% CO_2_ in a humidified incubator. Cells were plated in 384-well plates (Greiner Bio-One).

### Bacterial strains and plasmids. See Table 2

#### Bacterial Culture

*L. pneumophila* strain Paris wild-type and T4SS-defective Δ*dotA* mutant, both containing the pNT28 plasmid constitutively expressing GFP and a resistance cassette to chloramphenicol, were grown for 3 days on N-(2acetamido)-2-amino-ethanesulfonic acid (ACES)-buffered charcoal-yeast (BCYE) extract agar, at 37°C with 10µg/mL of chloramphenicol (SIGMA). On the day prior to the infection, 10 mL of Buffered Yeast Extract (BYE) liquid medium with 10µg/mL of chloramphenicol were inoculated with the bacteria at an optical density of OD_600_ = 0.1. The bacterial culture was allowed to grow overnight with shaking at 37°C and 200 rpm until the OD_600_ reaches approximately 4.2. The bacterial culture was diluted in the final cell culture medium to achieve a multiplicity of infection (MOI) of 10 for the subsequent experiments.

**Table 2.** List of bacterial strains and plasmids used in this study. * Channel (excitation/emmision, in nm): Green = 488/500-550. N/A = Non-Applicable.

| Species/Plasmid | Relevant genotype/description | Antibiotic Selection | Channel* | Ref. |
| --- | --- | --- | --- | --- |
| Lpp WT | Wild-type *Legionella pneumophila* Paris | - | N/A | (65) |
| Lpp ΔdotA | T4SS-defective *Legionella pneumophila* Paris | Kan<sup>r</sup> | N/A | (23) |
| Lpp ΔdotA | T4SS-defective *Legionella pneumophila* Paris | - | N/A | This study |
| Lpp WT DualFluor | *Legionella pneumophila* Paris with sfGFP under a synthetic constitutive promoter and sfmCherry under a *Ptet* promoter | Apr<sup>r</sup> | Green | This study |
| pNT28::GFP | pMMB207C, *egfp*, constitutive | Cam<sup>r</sup> | N/A | (45) |
| pNT28::sfmCherry | pMMB207C, *sfmCherry*, constitutive | Cam<sup>r</sup> | N/A | This study |
| pJM028 | pGEM AprR *mazF* | Apr<sup>r</sup> | N/A | This study |

#### Construction of bacterial strains

Natural transformation was employed to edit the genome of *L. pneumophila* Paris (66). Approximately 500 bp of sequence upstream and downstream of the gene interest was used to flank an apramycin resistance cassette and mazF gene taken from pJM028. Fragments were amplified by PCR (primers used in this study are shown in STAR Table) then ligated into pGEM T easy (PROMEGA) cut with HF-NotI (Thermofisher) using NEBuilder HiFi master mix (New England Biolabs). Resulting plasmids were transformed into E. coli DH5α and used as the template to generate approximately 1 µg of linear DNA by PCR, which was transformed as previously. Strains were checked by PCR and sequencing.

#### Infection of hMDMs

hMDMs were infected with *L. pneumophila* resuspended in the X-VIVO15 medium without Phenol Red (Lonza), at an MOI of 10, considering that an OD_600_ of 2.5 corresponds to 2.2 × 10^9^ bacteria/mL. To synchronize the infection, the plate was spun at 200 g for 5 minutes at room temperature. Following this, the plate was placed in a water-bath at 37°C for 5 minutes to promote phagocytosis of bacteria by the macrophages. Then, the plate was further incubated in a humidified incubator at 37°C with 5% CO_2_ for 25 minutes. After incubation, the cells were washed three times to remove extracellular bacteria.

#### Staining and confocal image acquisition

Specific fluorescent dyes were utilized to measure cellular parameters in macrophages during infection. For this, half of the culture medium was removed from the 384-well plate wells and dye mixes were added at 2X in a volume corresponding to the removed one, each at specific concentrations (see **Table 1**). The cells were incubated for 30 minutes at 37°C with 5% CO_2_ in a humidified incubator. Image acquisition was performed with an automated microlens-enhanced spinning disc confocal microscope (Opera Phenix High Content Screening System, PerkinElmer) with a 63x water objective. Excitation lasers were at 405, 488, 561, and 640 nm and emission filters were at 450, 540, 600, and 690 nm. Imaging of multiple fields (ranging from 9 to 25) was performed during 18 hours in an incubation chamber at 37°C with 5% CO_2_. Detailed protocol for imaging can be found at ref (46).

#### Image Analysis

Image analysis was performed with the Harmony software v.4.9 (Perkin Elmer) and self-built scripts by segmenting cell nuclei with Hoechst signal, cytoplasm with Cell Tracker Blue background, intracellular bacteria with GFP signal, and measuring metabolic parameters using red and/or far-red signals (**Table 1**). Open-source alternatives like CellProfiler (67), can also be used for this task. Output parameters for each cell included intensity and morphological measurements for each compartment and cellular structure. Our image analysis pipeline tracked the trajectories of every single cell over time using Harmony software (**Video S1**), but open-source solutions like TrackMate (68) can also be integrated to perform cell trajectory tracking. The data generated from image analysis was exported as a CSV dataset.

#### BATLI and Machine Learning (ML)

BATLI runs a front-end coded in HTML and Javascript assembled with a backend coded in Python, all coordinated by Flask software (https://github.com/pallets/flask). Whole BATLI code and example datasets can be downloaded from the BATLI GitHub repository (https://github.com/pescoll/BATLI). BATLI features a user-friendly interface accessible via any web browser, consisting of four modules used sequentially: ‘Cleaner’, ‘Loader’, ‘Viewer’, and ‘Backtracking’. The ‘Cleaner’ module imports, automatically cleans, and adapts the CSV dataset for further use. It can standardize datasets from different equipment and image analysis pipelines. The cleaned CSV file is then loaded into the ‘Loader’ module, which displays the dataset’s metadata and provides an experiment overview. The ‘Viewer’ module allows the selection of various parameters from the dataset, such as bacterial vacuole size (i.e. bacterial area = sum of the area occupied by GFP-expressing bacteria) or cell death (Annexin V intensity in the cytoplasm), to visualize all single-cell trajectories from the time-lapse, high-throughput imaging experiments. The ‘Backtracking’ module classifies trajectories based on defined cellular fates (e.g., intracellular bacterial replication) and thresholds, categorizing trajectories into two classes. This module performs retrospective analyses of each parameter measured within each class. The software outputs single-cell datasets organized by cell fate classes and selected parameters from the backtracking analysis. All results are downloadable as graphs and tables (CSV files, example in **Table S2**). For the ML approach, the scikit-learn library was used (https://scikit-learn.org/stable/) and in-house scripts were developed to train a Logistic Regression model. We used the confusion matrix, AUROC and precision-recall curves to measure the performance of the classification using prediction metrics. Whole ML code and example dataset can be also downloaded at BATLI GitHub repository.

## Supplementary Information

## Supplementary Figures

**Figure S1:**
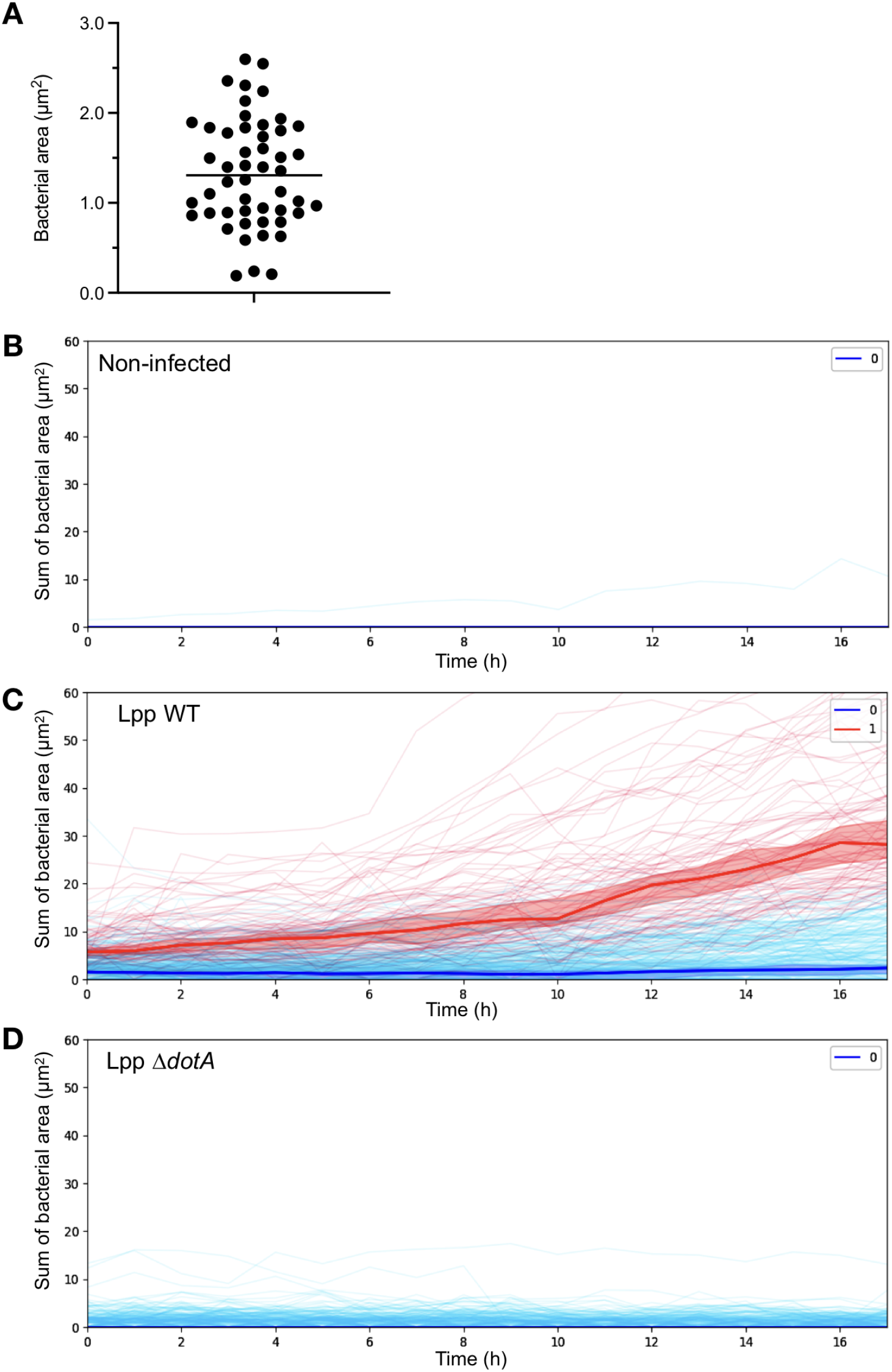
Related to Figure 1. **(A)** Distribution of bacterial area of single bacteria at t=0 hpi across 50 infected cells. The dot plot shows the area occupied by intracellular bacteria in individual cells measured in µm^2^. **(B)** Temporal progression of the sum of bacterial area in non-infected macrophages over 18 hpi. **(C)** Dynamics of bacterial replication in macrophages infected with *L. pneumophila* wild-type (Lpp WT). Individual cell trajectories are shown over time, with blue lines representing cells that did not support bacterial replication (Class 0), and red lines showing cells that supported bacterial replication (Class 1). **(D)** Temporal progression of bacterial area in macrophages infected with the avirulent *L. pneumophila* Δ*dotA* mutant strain. Blue lines represent non-replicating cells (Class 0), with no significant increase in bacterial area observed, confirming that the Δ*dotA* mutant strain is unable to replicate within macrophages.

**Figure S2:**
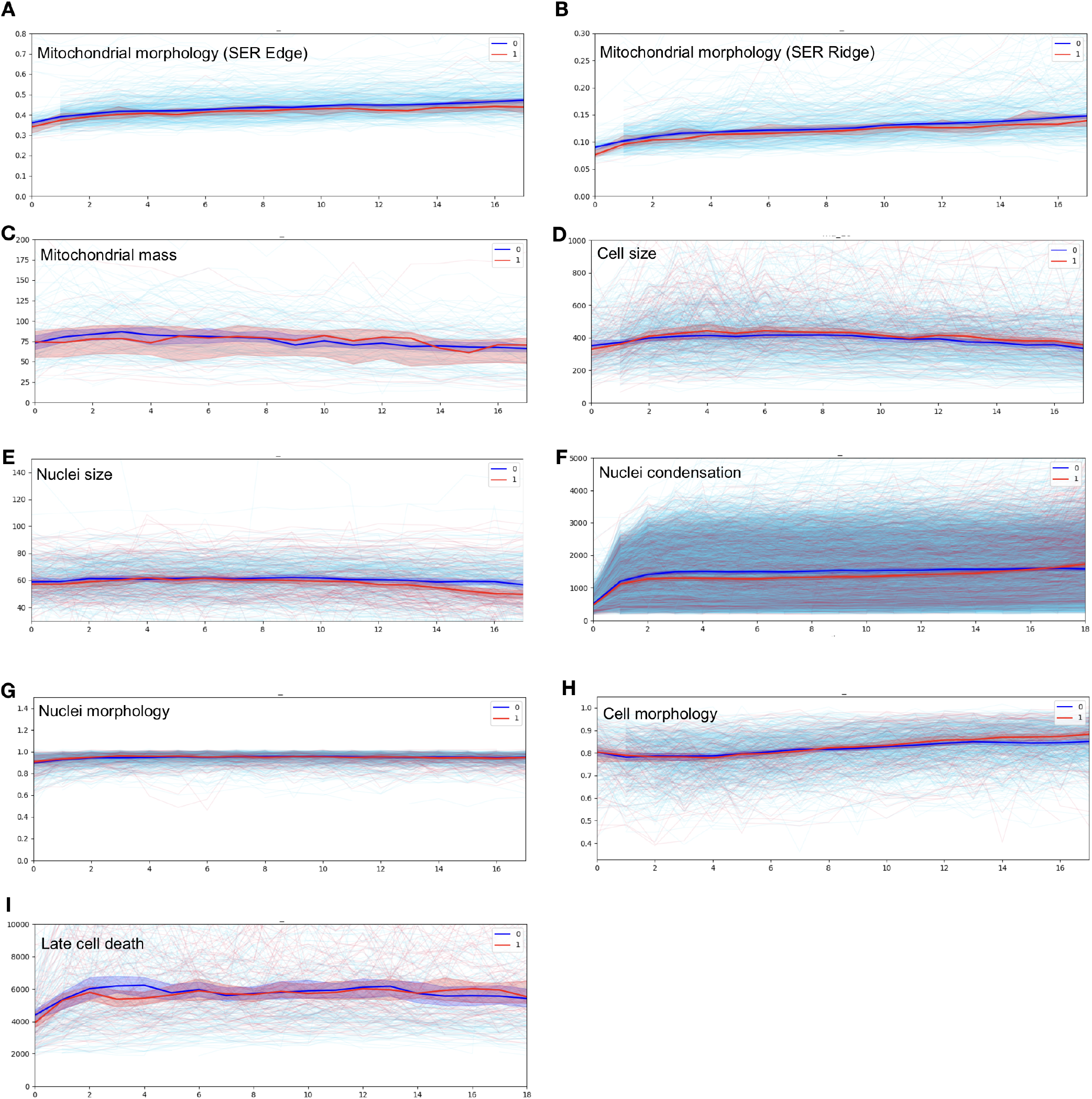
Related to Figure 2. BATLI analyses of various mitochondrial, nuclear, and cellular parameters in hMDMs infected with *L. pneumophila* wild-type (Lpp WT) over 18 hpi. The blue lines represent macrophages that did not support bacterial replication (Class 0), while the red lines represent macrophages that supported bacterial replication (Class 1). **(A)** Mitochondrial morphology using the SER Edge texture algorithm applied to Mitotracker signal (AU). **(B)** Mitochondrial morphology using the SER Ridge texture algorithm applied to Mitotracker signal (AU). **(C)** Mitochondrial mass as sum of the area of mitochondria (intracellular structures, spots) per cell using Mitotracker signal (µm^2^). **(D)** Cell Size (area of cytoplasm, µm2). **(E)** Nuclear size (area of nucleus, µm2). **(F)** Nuclei condensation (Intensity of Hoechst signal, AU). **(G)** Nuclear morphology (roundness of nucleus). **(H)** Cell morphology (roundness of cytoplasm). **(I)** Late Cell Death (SYTOX intensity, AU).

**Figure S3:**
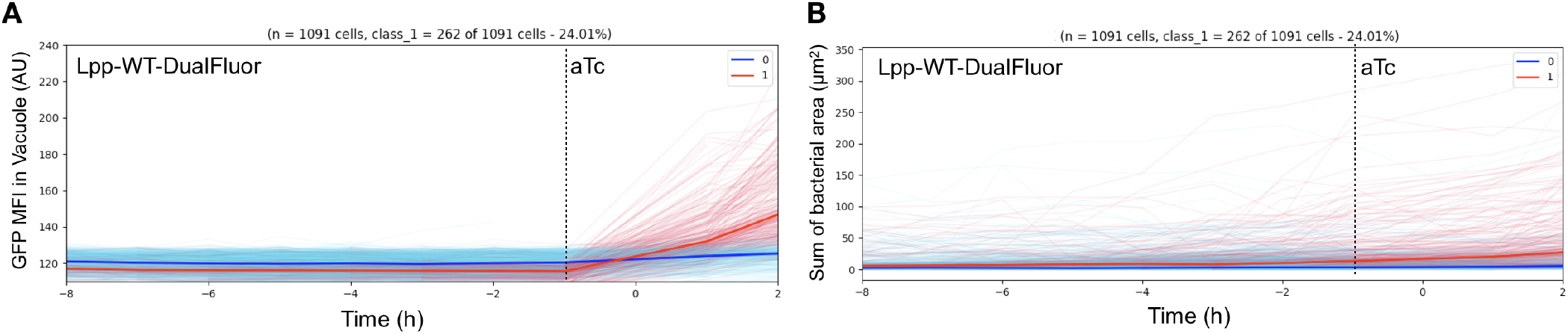
Related to Figure 4. **(A)** Classification of Lpp-WT-DualFluor intracellular bacteria according to the increase of GFP Mean Fluorescence Intensity (MFI). Blue lines (Class 0) are the trajectories of single bacterial vacuoles where GFP MFI did not increase after adding Anhydrotetracycline (aTc), while red lines shows bacteria where GFP MFI increased after adding aTc (Class 1). **(B)** Backtracking analysis of bacterial area (µm2) from the classification performed in (A).

**Figure S4:**
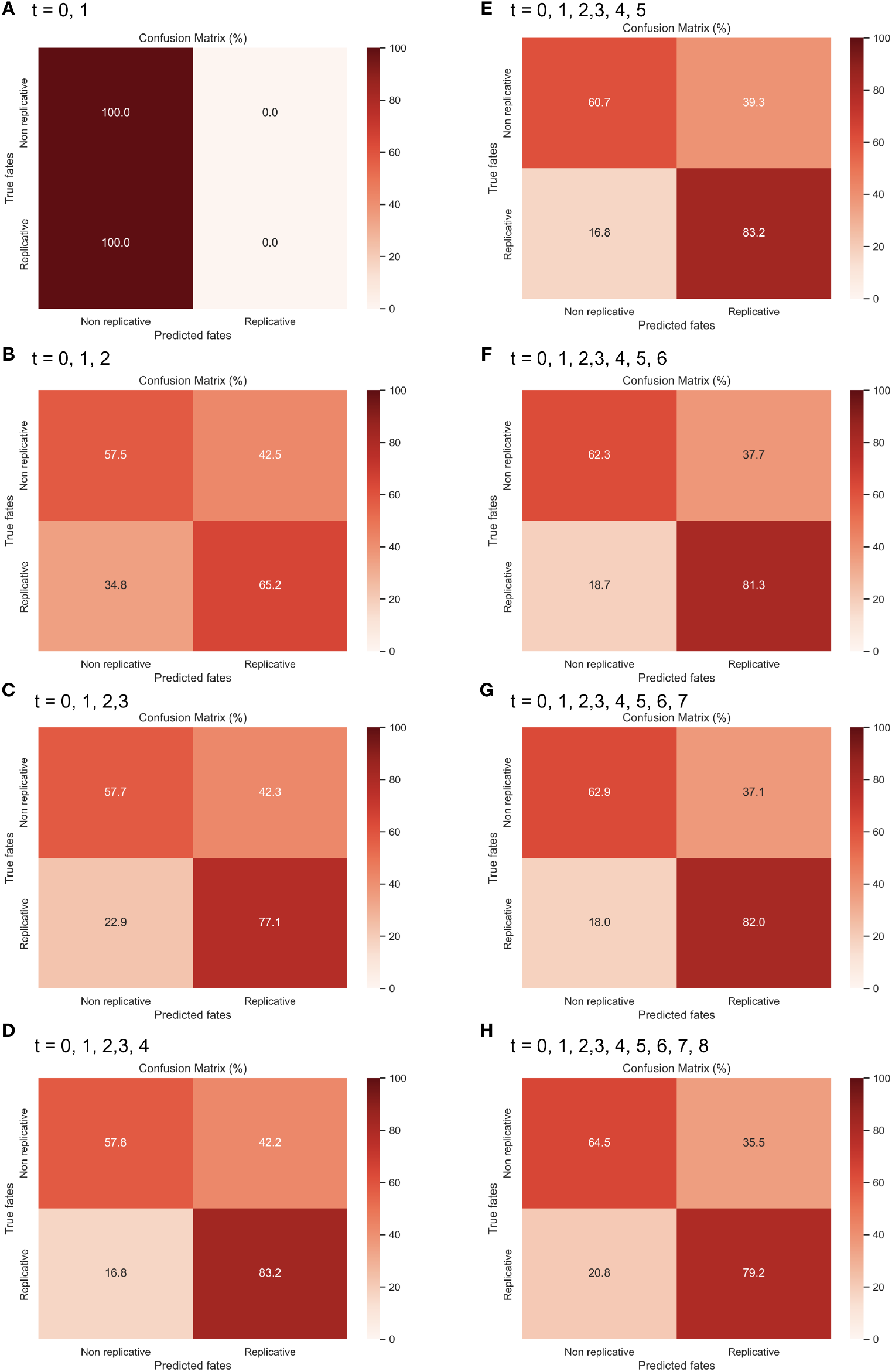
Related to Figure 5. Confusion matrices representing the performance of the logistic regression classification model in predicting whether human macrophages infected with *L. pneumophila* support bacterial replication or not, using the indicated ranges of times as training datasets. Each matrix compares the true cell fates (Y-axis) with the predicted fates (X-axis), measured as percentages. **(A)** 0 and 1 hpi, **(B)** 0, 1 and 2 hpi, **(C)** 0, 1, 2 and 3 hpi, **(D)** 0, 1, 2, 3 and 4 hpi, **(E)** 0, 1, 2, 3, 4 and 5 hpi, **(F)** 0, 1, 2, 3, 4, 5 and 6 hpi, **(G)** 0, 1, 2, 3, 4, 5, 6 and 7 hpi, **(H)** 0, 1, 2, 3, 4, 5, 6, 7 and 8 hpi.

## Supplementary Videos

**Video S1: Single-cell tracking of Lpp-WT infected, living hMDMs over time using Harmony software**. Time-lapse of confocal images from Lpp-WT-infected hMDMs. GFP-expressing Lpp-WT is shown in green. Hoechst-stained nuclei and Cell Tracker Blue-stained cytoplasm are shown in blue. Green and Blue signals were used to segment nuclear, cytoplasmic and bacterial compartments in each cell by automatic image analysis (white lines). Staining with TMRM dye to measure mitochondrial membrane potential (Δψm) is shown in yellow. The tracking of each cell is represented as white arrows showing the movement of the cell in the space. Some macrophages in the bottom of the video show intracellular replication of Lpp-WT.

## Supplementary Tables

**Table S1: Related to Figure 2**. Repeated-measures one-way ANOVA (Holm-Šídák’s multiple comparisons test) of the percentage of hMDMs where Lpp-WT bacterial area doubled with respect to the mean area of previous time points (minimum 2 consecutive time points). These results of significance test were used to generate Figure 2E.

**Table S2: Related to Figure S1**. Example of CSV table provided by BATLI containing all the parameters measured at the single-cell level and the corresponding classification label of each cell according to bacterial replication fate. Graphs from these analyses are those shown in Figures S1B-S1D.

